# Competition sensing alters antibiotic production in *Streptomyces*

**DOI:** 10.1101/2020.01.24.918557

**Authors:** Sanne Westhoff, Alexander Kloosterman, Stephan F. A. van Hoesel, Gilles P. van Wezel, Daniel E. Rozen

## Abstract

One of the most important ways that bacteria compete for resources and space is by producing antibiotics that inhibit competitors. Because antibiotic production is costly, the biosynthetic gene clusters coordinating their synthesis are under strict regulatory control and often require “elicitors” to induce expression, including cues from competing strains. Although these cues are common, they are not produced by all competitors and so the phenotypes causing induction remain unknown. By studying interactions between 24 antibiotic-producing streptomycetes we show that inhibition between competitors is common and occurs more frequently if strains are closely related. Next, we show that antibiotic production is more likely to be induced by cues from strains that are closely related or that share secondary metabolite biosynthetic gene clusters (BGCs). Unexpectedly, antibiotic production is less likely to be induced by competitors that inhibit the growth of a focal strain, indicating that cell damage is not a general cue for induction. In addition to induction, antibiotic production often decreased in the presence of a competitor, although this response was not associated with genetic relatedness or overlap in BGCs. Finally, we show that resource limitation increases the chance that antibiotic production declines during competion. Our results reveal the importance of social cues and resource availability in the dynamics of interference competition in streptomycetes.

**SIGNIFICANCE STATEMENT:** Bacteria secrete antibiotics to inhibit their competitors, but the presence of competitors can determine whether these toxins are produced. Here, we study the role of the competitive and resource environment on antibiotic production in *Streptomyces*, bacteria renowned for their production of antibiotics. We show that *Streptomyces* are more likely to produce antibiotics when grown with closely related competitors or that share biosynthetic pathways for secondary metabolites, but not when they are threatened by competitor’s toxins, in contrast to predictions of the competition sensing hypothesis. *Streptomyces* also often reduce their output of antibiotics when grown with competitors, especially under nutrient limitation. Our findings highlight that interactions between the social and resource environments strongly regulate antibiotic production in these medicinally important bacteria.

## INTRODUCTION

Bacteria live in diverse communities where they compete with other microbes for resources and space. Competition between different species can be regulated by the differential uptake and use of specific nutrients. It can also be driven by secreted toxins, like antibiotics or bacteriocins, that kill or inhibit competitors. Antibiotics and bacteriocins can allow producing strains to invade established habitats or repel invasion by other strains (1, 2). However, these compounds are expected to be metabolically expensive to make and so should only be produced against genuine threats from competitors. The competition sensing hypothesis predicts that microbes should upregulate toxin production when they experience cell damage or nutrient limitation caused by competitors (3). Alternatively, bacteria can also sense competitors by detecting secreted signals that predict imminent danger, but that cause no direct harm themselves, for example small molecule or peptide signals that are used to regulate toxin production by quorum sensing (3). Consistent with the predictions of competiton sensing, several studies have observed that microbes facultatively increase antibiotic or bacteriocin production when they are grown in co-culture with a competing strain (4–7). However, bacteria do not respond to all competitors in this way (7, 8).

Moveover, counter to predictions of competition sensing, co-cultivation can also cause strains to reduce antibiotic production (4, 6), rather than to respond aggressively to provocation. Why do bacteria respond to some competitors with aggression, but not to others? Similarly, are some cues from competitors more likely to elicit responses than others? To date, the answers to these questions have remained unknown. This shortcoming limits our ability to identify and induce cryptic antibiotic gene clusters for drug discovery and prevents a detailed understanding of the factors regulating the competitive dynamics of bacterial populations.

The aim of this paper is to address these issues in the context of bacteria from the prolific antibiotic-producing family *Streptomycetaceae* (9). These filamentous, spore-forming bacteria are renowned for their production of secondary metabolites, including many clinically useful antibiotics, anti-helminthic agents and anti-cancer drugs (10). Antibiotic production in Streptomycetes is associated with the developmental stage of the colony and typically coincides with the onset of sporulation (11, 12). We refer to this type of autonomous production as “constitutive” because it occurs in the absence of influence from other species. In addition, we and others have found that the presence of other strains in co-culture can alter antibiotic production by increasing or reducing antibiotic output (4, 5, 13). When they occur, these changes are thought to be caused by different social cues that indicate the presence of competitors. These can include nutrient stress, such as iron depletion, or factors that cause cellular damage or predict immediate danger, like antibiotics or quorum-dependent regulators of antibiotic production, such as gamma-butyrolactones (3, 14–16). However, as yet, we remain unable to predict the generality of these responses or the phenotypic or genomic factors that regulate them. Here, building on the framework of the competion sensing hypothesis, we set out to test if strains respond antagonistically to competitors that cause them harm. In addition, we investigate whether strains respond to competitive cues that we expect to be produced by strains with similar primary and secondary metabolism due to shared resource requirements or mechanisms of antibiotic regulation. Because these traits are phylogenetically conserved (6, 17–20), this predicts that *Streptomyces* will be more likely to respond to social cues from closely related species.

To examine the social factors that regulate antibiotic production in Streptomycetes, we studied antagonistic interactions between 24 different strains across a broad phylogenetic range in two nutrient environments. First, in each nutrient environment, we tested all possible pairwise interactions between these strains (24 × 24 = 576) by growing them as colonies and then testing if they could inhibit the growth of each other strain by inoculating these on top of the focal colony (Fig. 1). Next, we tested if growth in co-culture with a second strain altered the inhibitory behaviors we recorded during pairwise interactions. These three-way interactions (a total of 24 × 24 × 24 = 13,824 unique interactions in each nutrient environment) allowed us to compare the inhibitory capacity of strains during solitary growth, reflecting constitutive expression, to their behavior after interacting with a competitor during co-culture (Fig. 1). These approaches allowed us to directly test if altered antibiotic production during growth in co-culture could be predicted as a function of the phenotype or genotype of the competitor.

**Fig. 1.**
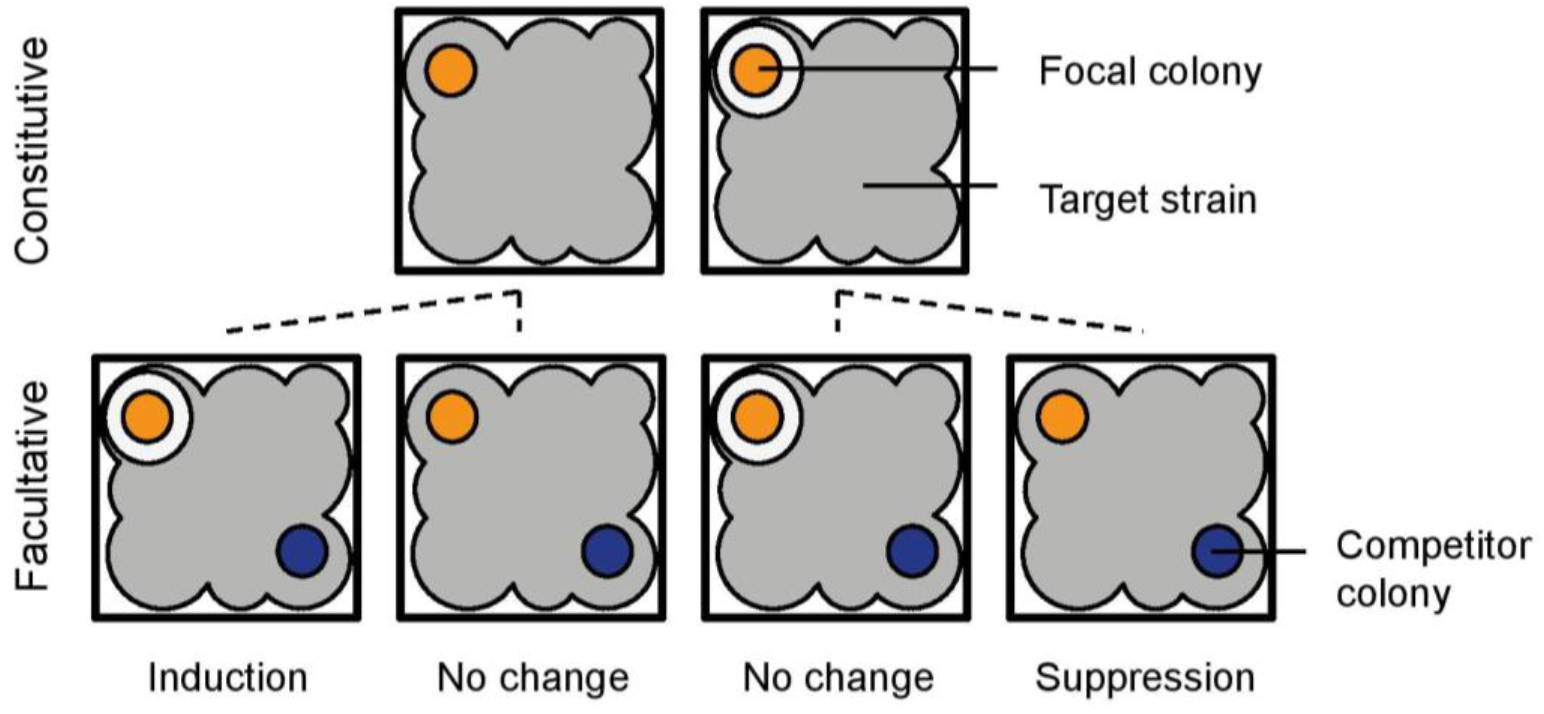
Schematic of constitutive and facultative inhibition assays. Focal strains (orange) were tested for their capacity to inhibit each target strain (grey) inoculated on top of the focal colony in a soft agar overlay. Inhibition was detected as a zone of clearance surrounding the colony. All 24 strains were tested as both focal and target strains, leading to 576 possible assays for constitutive antibiotic production. For the facultative assays a second colony was inoculated one centimeter away, designated as the competitor, that could interact with the focal strain through diffusible molecules. All 24 strains were tested as the focal, competitor and target strain, resulting in 24 × 24 × 24 = 13,824 assays. All assays were conducted in both high and low resource coditions. Comparing the ability of the focal strain to inhibit the target in the constitutive and facultative assays revealed whether antibiotic production was induced, suppressed or unchanged.

## RESULTS

### Constitutive antagonism

We first measured constitutive antibiotic production by growing each strain on a defined minimal medium and then testing if it could inhibit an overlay of each target strain (Fig. 1). These results formed the baseline against which we examined facultative responses. Similar to antagonistic interactions in *Streptomyces* and other microbes (4, 6, 21), these assays revealed that approximately half of all possible pairwise interactions were inhibitory (47.7%) (Fig. 2A, top left triangles). We used multilocus sequence typing (MLST) to infer the phylogeny of the strains. Next, we identified the biosynthetic gene clusters in the complete genomes of these strains using the bioinformatics tool antiSMASH (22). This revealed considerable variability in the number of secondary metabolite BGCswithin each genome (mean = 34 +/− 1.85 (SE), range = 22 to 64) (Fig. S2).

**Fig. 2.**
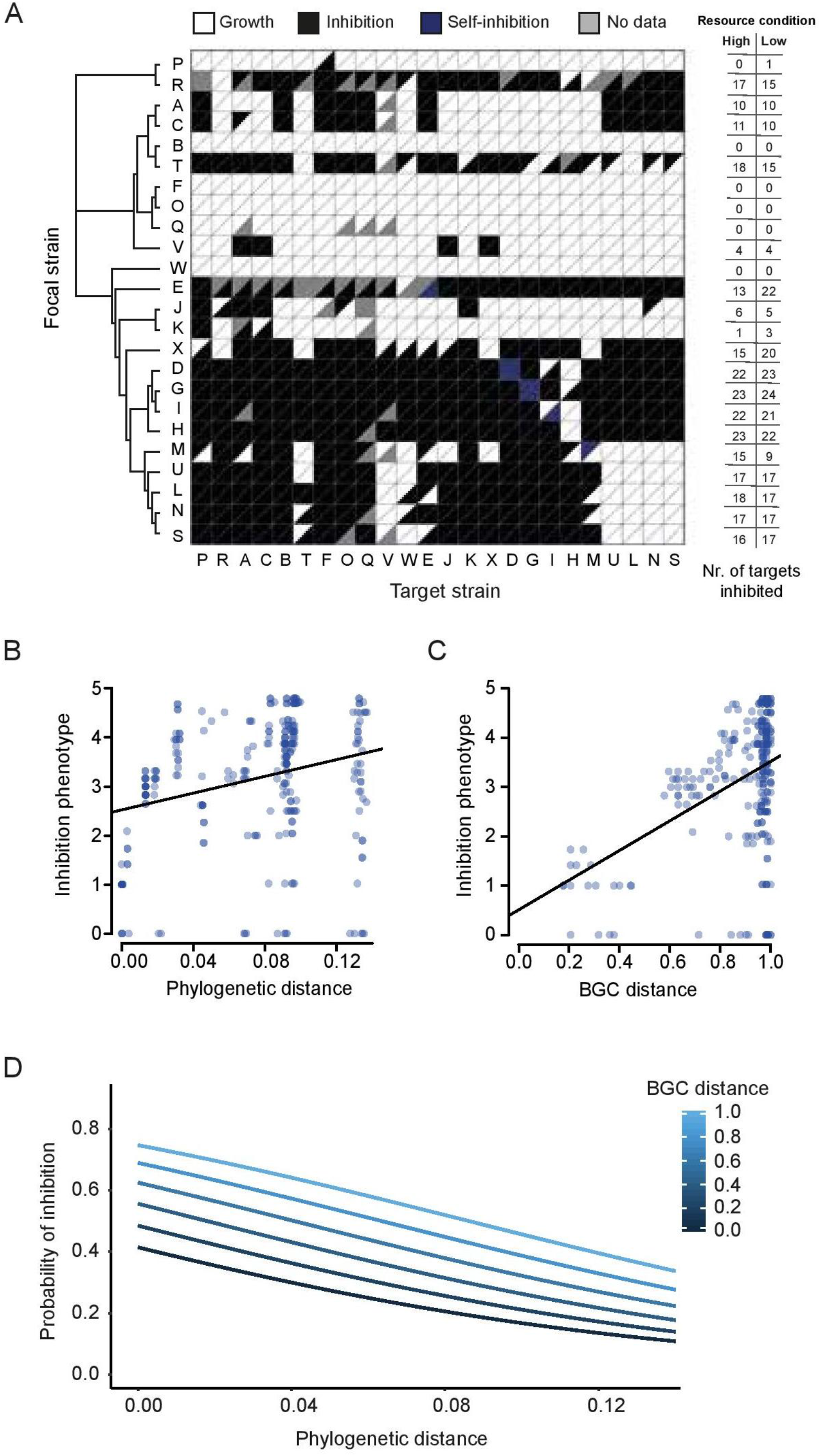
Constitutive antagonism. (A) Inhibition matrix sorted by multilocus sequence typing (MLST) relatedness. Triangles indicate whether a target strain showed growth (white) or was inhibited (black) by the focal strain. Each square is divided into two triangles: the upper triangle shows the results under high resource conditions, while the lower triangle shows the results under low resource conditions. Self-inhibition is denoted in blue. Missing data due to inconsistent results is shown in grey. B-D show results of assays conducted at high resource levels. (B) Correlation between inhibition phenotype dissimilarity (Euclidian distance determined by calculating the dissimilarity between focal strain inhibition phenotypes) and phylogenetic distance (Mantel test, *P* < 0.001, r = 0.27 N = 552) or (C) biosynthetic gene cluster (BGC) distance (Mantel test, *P* < 0.001, r = 0.43, N = 552). (D) Logistic regression between the probability of inhibition and phylogenetic and biosynthetic gene cluster (BGC) distance (*P*_phylogenetic distance_ < 0.001, *P*_BGC distance_ = 0.064, McFadden R^2^ = 0.02, N = 536).

The antagonistic behavior of each strain against the 24 possible targets generated a unique inhibition fingerprint, which we designate the inhibition phenotype. We calculated the dissimilarity in inhibition phenotypes to quantify whether strains inhibit the same or different targets and found a significant correlation between inhibition phenotype dissimilarity and phylogenetic distance (Fig. 2B) (Mantel test, *P* < 0.001, r = 0.27). This result indicates that closely related strains inhibit the same targets. We then tested if this was due to the possibility that related strains produce similar antagonistic compounds. To address this, we grouped all BGCs identified in the genomes into gene cluster families using BiG-SCAPE (23) and calculated Jaccard distances between the strains based on their shared BGCs. This analysis revealed that BGC distance is significantly correlated with inhibition phenotype (Mantel test, *P* < 0.001, r = 0.43) (Fig. 2C), supporting the idea that related strains produce similar secondary metabolites.

Consistent with the idea that closely related strains are more likely competitors, strains showed a stronger tendency to inhibit closely related targets (logistic regression, *P* < 0.001, McFadden R^2^ = 0.02, N = 536). As BGCs often also provide resistance against the product they encode, we expected that strains with a high degree of BGC similarity would not inhibit each other. Indeed, strains were most likely to inhibit targets that are closely related but have dissimilar BGCs (logistic regression, *P*_phylogenetic distance_ < 0.001, *P*_BGC distance_ = 0.064, McFadden R^2^ = 0.02, N = 536) (Fig. 2D). In contrast to another study that examined inhibitory interactions between phylogenetically diverse bacteria (21), we found no association between the probability of inhibition and the metabolic overlap between strains, assessed as the ability to grow on 95 different carbon sources using BiOLOG plates.

### Altered inhibition during co-culture

Our results show that Streptomycetes constitutively produce antibiotics that inhibit closely related strains. However, constitutive antibiotic production does not account for facultative changes that are caused by cues from other strains. We measured facultative responses by inoculating each strain next to a competitor and then assessing if it could inhibit the growth of the different target strains, as above. This allowed us to directly compare the inhibitory capacity of each focal strain in the presence and absence of each competitor (Fig. 1). A focal-competitor interaction was scored as induced if the focal strain was able to inhibit any of the target strains that it was unable to inhibit when it was grown on its own, and suppressed if the opposite occurred. By this approach, a focal strain could be both induced and suppressed by the same competitor, against different target strains. The results of these assays, shown in Figure 3A, confirm that facultative responses are extremely widespread. The inhibition phenotype was changed by a competitor in approximately half of the focal-competitor interactions (48%), meaning that the focal strain was induced or suppressed against at least one target strain in the presence of a given competitor. These changes dramatically altered the inhibition phenotype of the focal strain and changed the total number of strains that each focal strain could inhibit (Fig. 3B). Overall, we observed induction in 33% of all tested focal-competitor interactions and suppression in 45%. There was considerable variability in the responsiveness of strains to competitors; whereas some strains responded to none of the competitors, others responded to nearly all of them (induced: 0-20, suppressed: 0-22) (Fig. 3C). On average each focal strain was induced by 7.4+/− 1.5 (SE) competitors and suppressed by 9.6 +/− 1.7 (SE). In many cases, a given strain was both induced and suppressed by the same competitor against different targets (Fig. 3A). Although this led to a distinct inhibition phenotype, as compared to the focal strain grown alone, it may not have changed the total number of inhibited targets.

**Fig. 3.**
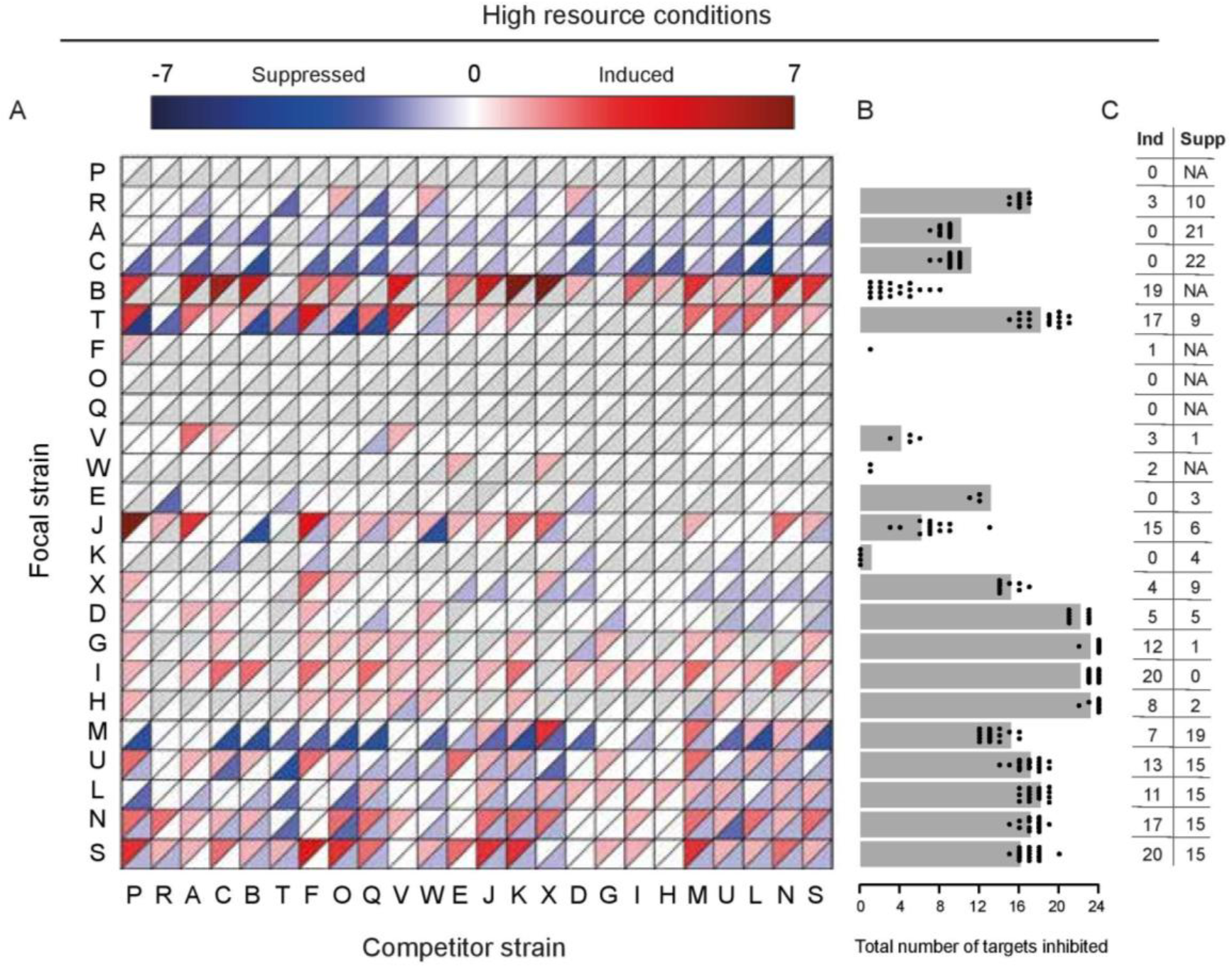
Altered antagonism during co-culture under high resource conditions. (A) Interaction heatmap showing changes to target strain inhibition when a focal strain is grown in co-culture with a competitor. Induction is shown in red, while suppression is shown in blue. Grey triangles indicate that either induction or suppression was not possible for this focal strain. (B) Grey bars indicate the number of target strains inhibited by the focal strain when grown alone. Black dots indicate the net number of target strains inhibited by the same focal strain if it was induced and/or suppressed during co-culture with one of the 24 possible competitors. Dots showing the same number of inhibited target strains as the grey bar indicate that a competitor strain causes an equal level of induction and suppression against targets, resulting in no net change. (C) Number of competitors that induce or suppress each focal strain. Cases where suppression is not possible due to the absence of constitutive inhibition are denoted as NA.

Competition sensing predicts that bacteria will change their behavior in response to antagonistic competitors that they detect by sensing cell damage (3). We define a competitor as antagonistic if it inhibits the focal strain during the constitutive assay. Although we found that induction of the focal strain was significantly related to whether or not its competitor was antagonistic (logistic regression, *P* < 0.001, McFadden R^2^ = 0.06, N = 354), the direction of this result did not match our expectations (Fig. 4A, black bars). Unexpectedly, antibiotic production in the focal strain was nearly twice as likely to be induced by a non-antagonistic competitor (probability of induction 0.41 vs 0.22). This indicates that cell damage was not a strong cue for antibiotic induction. Other ways that focal cells could sense competitors is if they detect compounds they produce, such as antibiotics and quorum sensing signals, or by experiencing nutrient stress due to resource competition. Since both primary and secondary metabolism are correlated with phylogenetic distance, we examined if induction was correlated with phylogenetic distance. As predicted, focal strains are more frequently induced by a closely related competitor (logistic regression, *P* < 0.001, McFadden R^2^ = 0.02, N = 487) (Fig. 4B, solid line). To examine if this effect was driven by the production of similar secondary metabolites, we tested if differences in induction could be explained by BGC similarity. Indeed focal strains are more likely induced by competitors with which they share more BGCs (logistic regression, *P* < 0.001, McFadden R^2^ = 0.04, N = 487) (Fig. 4C, solid line). This suggests that cues for induction rely more on the detection of secreted compounds, rather than the damage these compounds cause.

**Fig. 4.**
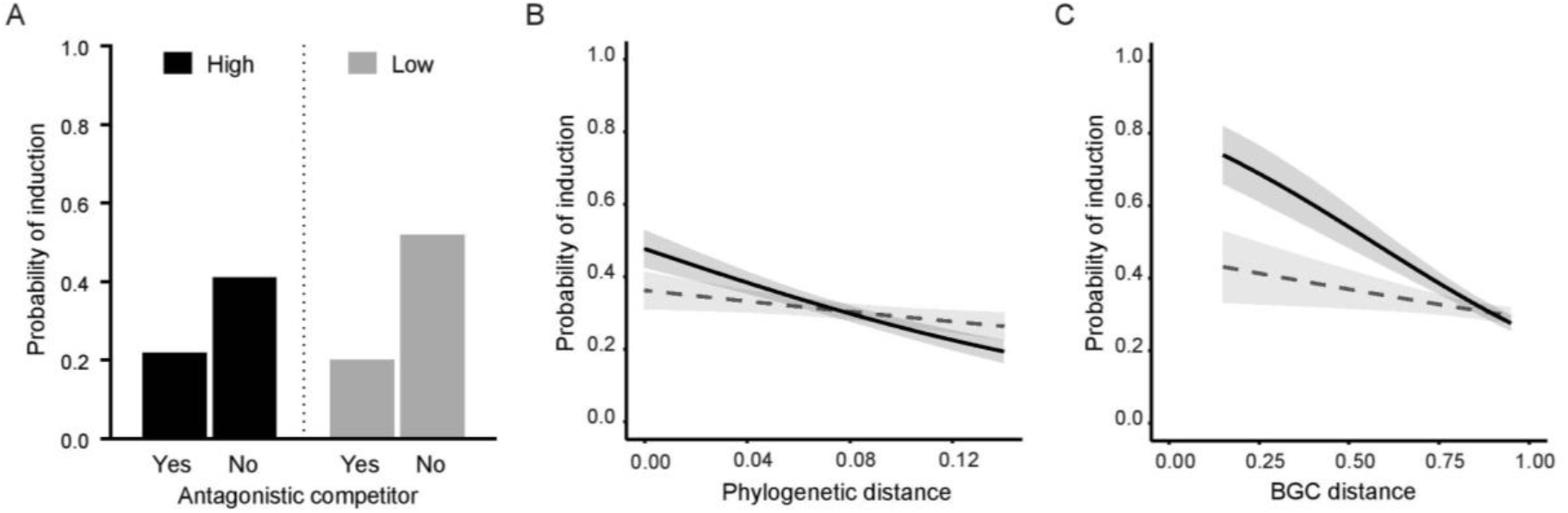
Induction during co-culture. (A) The probability that a focal strain is induced is lower when the competitor is antagonistic to the focal strain under both high (Logistic regression, *P* < 0.001, McFadden R^2^ = 0.06, N = 354) and low resource conditions (Logistic regression, *P* < 0.001, McFadden R^2^ = 0.12, N = 419). (B) Logistic regressions between the probability of induction and phylogenetic distance under high (black line) and low resource conditions (dashed line) (*P* < 0.001, McFadden R^2^ = 0.02, N = 487 and (logistic regression, *P* = 0.205, McFadden R^2^ = 0.00292, N = 445 respectively) or (C) Logistic regressions between the probability of induction and BGC distance under high (black line) and low resource conditions (dashed line) (*P* < 0.001, McFadden R^2^ = 0.04, N = 487 and *P* = 0.166, McFadden R^2^ = 0.00341, N = 445 respectively). Ribbons indicate SE.

In addition to induction, we found that antibiotic production was also commonly suppressed in the presence of competitors. Although this strategy can be perceived as benefiting the competitor strain if it prevents a focal strain from producing a potentially harmful antibiotic, it could also benefit the suppressed strain by allowing it to redirect energy towards other functions. However, we found no relationship between suppression and the competitor’s ability to inhibit the focal strain (logistic regression, *P* = 0.83, McFadden R^2^ = 0.025). Suppression was also not associated with phylogenetic or BGC distance (logistic regression, *P* = 0.94, McFadden R^2^ = 0.0000123, N = 366 and *P* = 0.166, McFadden R^2^ = 0.00384, N = 366 respectively).

### Effect of resource stress on inhibition

To address the role of nutrient limitation on antibiotic production, we tested whether constitutive or facultative inhibition changed if the carbon source concentration was reduced by 10-fold (Fig. 2A, bottom right triangles). The frequency of constitutive inhibition was marginally higher under these conditions: 49.2% vs 47.7% of all pairwise interactions were inhibitory on low versus high resource medium, and only 6.7% of pairwise interactions differed between the two resource conditions (McNemar’s Χ^2^ = 0, df = 1, *P* = 1). Likewise, we found a strong correlation between the inhibition phenotypes of the strains at both resource concentrations (Mantel test, r = 0.93, *P* < 0.001), with phylogenetic and BGC distance both significantly correlated with inhibition phenotype, and in the same direction as in the high resource concentration (Mantel test, *P* < 0.001, r = 0.30 and *P* < 0.001, r = 0.39 respectively) (Fig. S1). As at the higher glycerol concentration, strains are more likely to inhibit closely related targets with dissimilar BGCs (logistic regression, *P*_phylogenetic distance_ < 0.001, *P*_BGC distance_ = 0.023, McFadden R^2^ = 0.02, N = 526) (Fig. S1).

Streptomycete focal strains responded differently to the presence of a competitor under varying resource conditions (Fig. 5) (McNemar’s Χ^2^ = 5.43, df = 1, *P* < 0.05), with a change in inhibition phenotype in 56.1% versus 49.1% in low versus high resource conditions, respectively (Fig. 6). While we expected more induction due to resource stress in accordance with the competition sensing hypothesis, the incidence of induction was slightly lower at lower resource levels (30.2% vs 33.0%). By contrast we observed a dramatic increase in suppression, from 45.0% to 59.1% (Fig. 6). Just as at the higher resource level, strains were more likely to be induced by competitors that did not inhibit them (Fig. 4A, grey bars) (Logistic regression, *P* < 0.001, McFadden R^2^ = 0.12, N = 419). In contrast to the higher resource conditions, neither phylogenetic nor BGC similarity was associated with induction in the low resource environment (Fig. 4B and C, dashed lines) (logistic regression, *P* = 0.205, McFadden R^2^ = 0.00292, N = 445 and *P* = 0.166, McFadden R^2^ = 0.00341, N = 445 respectively). Suppression was still not associated with any of the factors that we tested, suggesting that antibiotic suppression may be a general reaction to resource stress in Streptomycetes.

**Fig. 5.**
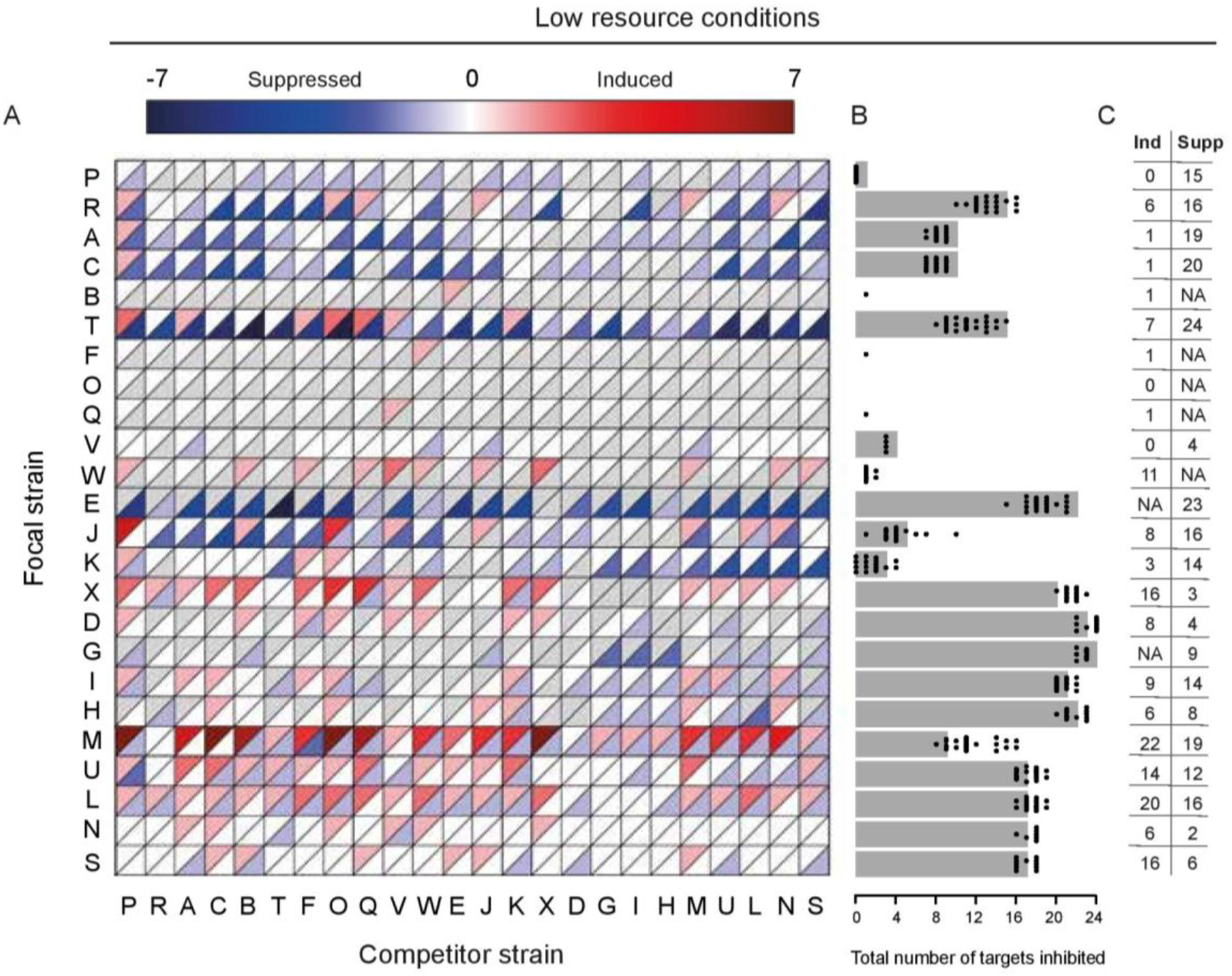
Altered antagonism during co-culture under low resource conditions. (A) Interaction heatmap showing change in target strain inhibition when a focal strain is grown in co-culture with a competitor. Induction is shown in red, while suppression is shown in blue. Grey triangles indicate that either induction or suppression was not possible for this focal strain. (B) Grey bars indicate the number of strains inhibited by the focal strain when grown alone. Black dots indicate the net number of target strains inhibited by the same focal strain if it was induced and/or suppressed during co-culture with one of the 24 possible competitors. Dots showing the same number of inhibited target strains as the grey bar indicate that a competitor strain causes an equal level of induction and suppression against targets, resulting in no net change. (C) Number of competitors that induce or suppress each focal strain. Cases where suppression is not possible due to the absence of constitutive inhibition are denoted as NA.

**Fig. 6.**
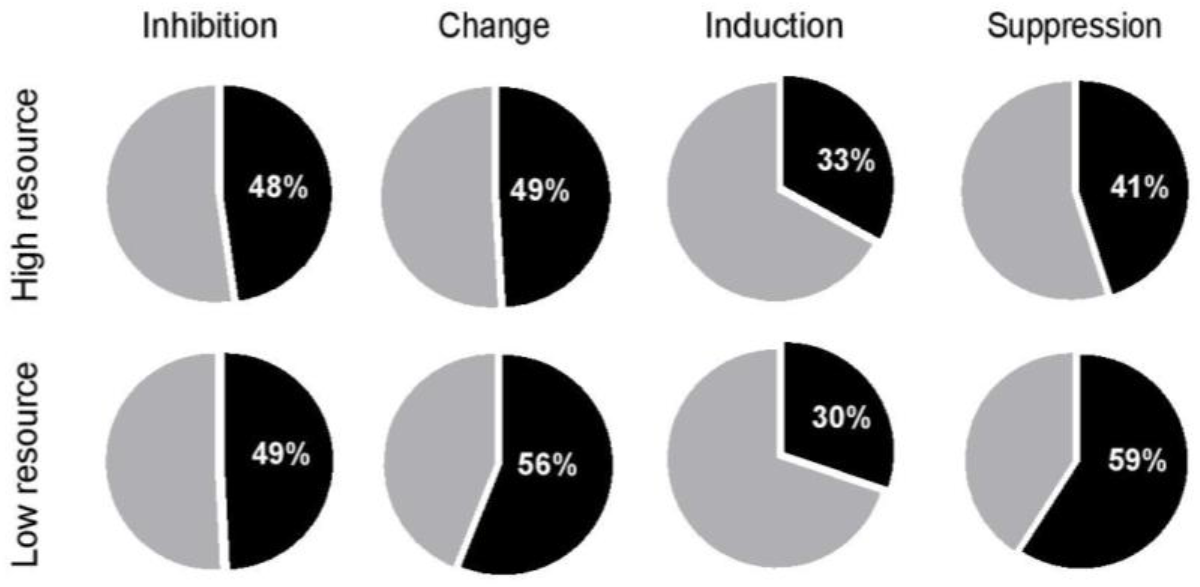
Constitutive and facultative inhibition under high and low resource conditions. Comparison of the total amount of inhibition, change in inhibition due to competition, induction and suppression found in low and high resource conditions.

## DISCUSSION

Competitive and social interactions between neighboring microbial cells in soil are common as different species vie for space and resources. One of the ways that bacteria compete is by secreting toxins like antibiotics or bacteriocins, but the production of these compounds is typically studied without consideration of this biotic context. Both theory and experiments have shown that this perspective is limited because it neglects factors that induce or suppress toxins and fails to identify toxins whose production is dependent on competitive interactions (4, 5, 24–27). In this context, the aims of our work were twofold: first to establish to what extent the role of social interactions affect antibiotic production in common soil microbes of the *Streptomycetaceae*, and second to identify factors that are predictive of competition-mediated responses.

By comparing antibiotic production in the absence and presence of another species in co-culture, termed constitutive and facultative production, respectively, we found that production was induced in ~ 1/3 of co-cultures, both in high and low resource conditions. These results considerably expand on results from earlier studies that describe changes in secondary metabolite production and inhibition in a smaller set of Actinomycetes or against a smaller set of target species (4–6). The latter aspect is especially important because susceptibility varies markedy between strains; studies with fewer targets may therefore underestimate the frequency of facultative changes to antibiotic production. Induction was strongly predicted by phylogenetic distance. A similar result in an earlier study was linked to nutrient use as closely related strains compete for the same resources, but did not test for overlap in secondary metabolites (6). This same study found that Streptomycetes increase antibiotic production when grown with susceptible competitors, without assessing the impact of whether or not the competitor could reciprocally inhibit the focal strain. Unexpectedly, we found that a focal strain was less likely to be induced during co-culture if it was grown with an inhibitory competitor. In other words, while co-culture frequently altered antibiotic production, this was not evidently driven by cellular damage caused by the second strain, as specifically predicted by the competition sensing hypothesis (3). Instead, our results suggest that cells are more likely to induce antibiotic production in response to cues that are correlated with phylogenetic distance, rather than direct harm itself. For example, under high resource conditions strains that share BGCs are more likely to induce each other. This suggests two possible sources for cues. First, antibiotic intermediates or antibiotics themselves, can serve as inducers of antibiotic production or resistance (16). These responses can prevent autotoxicity or killing by neighboring clonemates and also act as regulators of the expression of their own biosynthetic gene cluster (26, 28). Because resistance is often encoded in the antibiotic biosynthetic gene cluster, self-inhibition, which was observed in few instances, could be due to not-yet-expressed resistance in spores during challenge with a high dose of antibiotic, or alternatively by the production of germination inhibitors, such as germicidin, that inhibit the germination of conspecific spores (29). Second, related strains that share one or more BGCs may be more likely to utilize the same, or similar, secreted factors that induce antibiotic production, e.g. the quorum-dependent gamma-butyrolactone signals. *Streptomyces* contain multiple receptors for cognate and non-cognate gamma-butyrolactones, thereby allowing them to detect these signals as a precursor of the antibiotics another strain might produce (15, 30, 31). Similar eavesdropping of quorum-dependent signals has been observed for bacteriocins in *Streptococcus pneumoniae*, which leads to cross-induction of strain-specific antimicrobials (7). Testing this idea in *Streptomyces* using chemically synthesized signals and reporter strains remains an important objective for future work.

When we repeated our assays at 1/10 the glycerol concentration, constitutive expression was only marginally changed; however, these lower resource concentrations led to slightly reduced induction rates and a marked increase in suppression. Moreover, the associations between induction and phylogenetic distance and BGC distance disappeared. These results indicate that antibiotic regulation integrates information about the competitive environment as well as environmental resource availability, leading strains to respond differently when exposed to a combination of competitive cues and resource stress then when exposed to only one of these. Links between nutrient sensing or carbon catabolite repression and antibiotic production in Streptomycetes are well established. For example, under nutrient rich conditions N-acetylglucosamine blocks morphogenesis and antibiotic production, while it has the opposite effect under nutrient poor conditions (32, 33). A second possibility may be that competition exacerbates nutrient stress overall, leading to a general suppressive response that doesn’t depend on the particulars of the competitor. By this view, suppression is best considered as a generic response to nutrient stress, rather than the result of a specific action by the second strain. This result indicates that further work will need to consider responses other than antibiotic production when examining the behavior of cells in co-culture. For example, strains may respond to nutrient stress from competitors by redirecting energy used for antagonism towards functions that help them to avoid competition, e.g. hyphal growth in the direction opposite the competing strain or increased sporulation. Whereas the first possibility would contribute to an escape in space, the latter would allow an escape in time, leaving spores to germinate when nutrient stress is relieved. These alternative responses, which are equivalent to a “fight or flight” decision, might be anticipated if there are trade-offs between antibiotic production and other aspects of development, as we have found in *S. coelicolor* (34).

In summary, our results provide strong evidence that antibiotic production by streptomycetes is highly responsive to their social and resource environment. We establish the importance of BGC similarity on antibiotic induction, suggesting a role for shared regulatory compounds, and show that suppression and possibly escape, as a means of “flight”, should be more thoroughly examined as a response to interference competition.

This is equally important for many of the other mechanisms that bacteria use to regulate inter- and intra-specific warfare (35). It will also be crucial to examine these responses in experiments that more closely approximate the natural environment, including environments with increased spatial heterogeneity and decreased diffusion, and where local interactions are maintained over longer periods of time. Similarly, an important next step is determine how these social interactions influence competitive outcomes, as has been done for constitutive antibiotic production between competing species (1, 2, 36). Together, these approaches will lead to a fuller understanding of the role of antibiotic production in natural soils and the factors that maintain microbial diversity. In addition, they will help to identify factors that can be used to induce cryptic antibiotic BGCs in *Streptomyces* as potential drug leads.

## METHODS

### Strains and culturing conditions

The panel of 24 *Streptomycetaceae* strains used in this study (Table S1) included 21 strains isolated from a single soil sample from the Himalaya Mountains collected at 5000 m near a hot water spring (37). These 21 strains were selected due to their consistent phenotypes and the ability to sporulate in our lab growth conditions. The remaining three strains were well-characterized lab strains, *Streptomyces coelicolor* A3(2) M145, *Streptomyces griseus* IFO13350 and *Streptomyces venezuelae* ATCC 10712. High density spore stocks were generated by culturing on Soy Flour Mannitol Agar (SFM) (20 g Soy Flour, 20 g Mannitol, 20 g Agar per liter) or on R5 Agar (103 g sucrose, 0.42 g K_2_SO_4_, 10.1 g MgCl_2_, 50 g glucose, 0.1 g CAS amino acids, 5 g yeast extract, 5,7 g TES, 2 ml R5 trace element solution and 22 g agar per liter). After 3-4 days of growth, spores were harvested with a cotton disc soaked in 3 ml 20% glycerol, and spores were extracted from the cotton by passing the liquid through an 18g syringe to remove the vegetative mycelium. Resulting spore stocks were titred and stored at −20 °C.

Multi-well masterplates were prepared by diluting the high density spore stocks to 1 × 10^6^ sp ml^−1^ in deionized water and these plates were stored at −20 °C. The glycerol concentration after the dilution of stocks was always lower than the concentration of glycerol added as a carbon source to the medium.

To perform the interaction assays approximately 1 μl of the focal strain, and when indicated 1 μl of the competitor strain, was replicated on a 25 grid plate (Thermo Fisher Scientific, Newport, UK) using a custom built multi-pin replicator (EnzyScreen BV, Heemstede, The Netherlands) from a frozen masterplate. Each well of the 25 grid plate contained 2 ml Minimal Medium (MM) (500 mg L-Asparagine (Duchefa Biochemie, The Netherlands), 500 mg KH_2_PO_4_ (Duchefa Biochemie, The Netherlands), 200 mg MgSO_4_.7H_2_O (Duchefa Biochemie, The Netherlands), 10 mg Fe_2_SO_4_.7H_2_O (Sigma Aldrich, MO, USA) and 20 g agar (Company) per litre, pH 7.2 supplemented with either 0.05% or 0.5% glycerol). After 4 days of growth at 30 °C a 1 ml overlay (0.8% agar MM) containing 1.6 × 10^5^ sp/ml was added on top. After 24 to 48 hours of incubation at 30 °C (depending on the growth speed of target strain) 1 ml of the dye resazurin (Cayman Chemical Company, Michigan, USA) was added to each well at a concentration of 50 mg L^−1^ and incubated for half an hour before the surplus was removed. Change in colour of this redox dye from blue to pink was used as a measure of growth of the target strain, as resazurin (blue) is changed to resorufin (pink) by metabolically active cells. Pictures were taken of every plate and these were scored for the presence or absence of inhibition zones around the colony/colonies. Every interaction was assessed in duplicate. When the results of assays were inconsistent, the particular interaction was repeated a third time.

### Whole genome sequencing

Whole genome sequencing was performed for all strains for which a full genome sequence was not yet available to perform genome mining and to generate a phylogenetic tree. As described before (38) strains were grown in liquid culture containing 50% YEME/50% TSBS with 5mM MgCl_2_ and 0.5% glycine at 30 °C, 250 rpm for 2 days. After centrifugation the pellet was resuspended in TEG-buffer with 1.5 mg ml^−1^ lysozyme and after 1 hour of incubation at 30 °C the reaction was stopped by adding 0.5 volume of 2M NaCl. DNA was extracted using a standard phenol/chloroform extraction, followed by DNA precipitation and washing in isopropanol and 96% ethanol. Dried DNA was resuspended in MQ water and then treated with 50 ug ml^−1^ of RNase and incubated at 37 °C for 1 hour. Following RNase treatment, the mixture was purified and cleaned as above, after which the purified DNA was washed with 70% ethanol and resuspended in MQ water. Paired-end sequence reads were generated using the Illumina HiSeq2500 system at BaseClear. *De novo* assembly was performed using the “De novo assembly” option of the CLC Genomics Workbench version 9.5.1 and the genome was annotated using the BaseClear annotation pipeline based on the Prokka Prokaryotic Genome Annotation System (version 1.6).

Using the complete genomes, multilocus sequence typing (MLST) was performed as described by (39). For this purpose we used the sequences of six housekeeping genes, *atpD, gyrB, recA, rpoB, trpB* and 16S rRNA that were shown to give good resolution for the *S. griseus* glade. For the already available sequenced genomes, the sequences for *S. coelicolor* (strain V) were downloaded from StrepDB (http://strepdb.streptomyces.org.uk) and used to blast against the genome sequences of *S. venezuelae* ATCC 10712 (txid 54571) (strain W), *S. griseus* supsp. griseus NBRC 13350 (txid 455632) and MBT66 (strain P) on the NCBI database.

For all sequenced genomes the genes of interest were located from the annotated genome or were searched in a database constructed with the genomes in Geneious (Geneious 9.1.4). Each gene was aligned and trimmed before the six sequences for each strain were concatenated in frame and used to construct a neighbourjoining tree using Geneious to reveal the phylogenetic distances between the strains.

### Analysis of biosynthetic gene clusters

Biosynthetic gene clusters were identified within each genome with antiSMASH version 4.0 (22). BiG-SCAPE was used to calculate the pairwise distances between all BGCs, using a cutoff of 0.5 as a threshold for similarity (23). This generated a BGC presence/absence matrix that we used to calculate a Jaccard distance between each pair of genomes to define the BGC distance between the strains.

### Resource use

Carbon source utilization of each strain was tested using BiOLOG SFP2 plates (Biolog, Hayward, CA, USA) on which growth on 95 carbon sources can be assessed. Plates were inoculated as described by (40). Briefly, strains were grown on MM with 0.5% glycerol for 7 days before spores were swabbed into a 0.2% carrageenan solution and adjusted to OD_590_ of 0.2 – 0.24. This solution was diluted 10 times in 0.2% carrageenan and 100 ul of this dilution was added to each well. Plates were incubated at 30 °C for 3 days before the absorbance of each well at 590 nm was measured using a Spark 10M plate reader (Tecan, Switzerland). All strains were assessed in triplicate. For the analysis the absorbance of the water control was subtracted for each well and the average was taken. If the average was not significantly different from 0 (one sample T-test), the value was adjusted to 0. The Pearson correlation coefficient was calculated between all possible pairwise combinations of the strains and the metabolic distance was calculated as 1 – correlation coefficient. Strain P showed extremely poor growth on the BiOLOG plates and was therefore excluded.

### Statistics

All statistics were performed in R. Correlation between phylogenetic distance, metabolic dissimilarity, secondary metabolite distance and inhibition and resistance phenotype was determined using Mantel tests. To establish whether antagonism and inhibition, induction and suppression are dependent, logistic regressions were performed. Logistic regression was also used to test for association between inhibition, induction or suppression and phylogenetic distance, metabolic distance or BGC distance. For the logistic regressions we excluded all self-self interactions, as these confound the analyses by having zero distance between the strains or test for self-inhibition.

## ACKNOWLEDGEMENTS

This work was financially supported by a grant from the Dutch National Science Foundation (NWO) to DER. The authors contributed in the following ways: SW and DER designed the experiments; SW performed the experiments and analyzed the data; AK and SFAvH assisted with the (bio)informatics; and SW, DER and GPvW wrote the manuscript. The authors declare there is no conflict of interest.

