## Supplementary Material for "Competition sensing alters antibiotic production in *Streptomyces*"

Daniel E. Rozen*

Institute of Biology, Leiden University, Sylviusweg 72, 2333 BE Leiden, The Netherlands

* Corresponding authors: Sanne Westhoff and Daniel E. Rozen

**This PDF file includes:**

Figures S1 – S3

Table S1

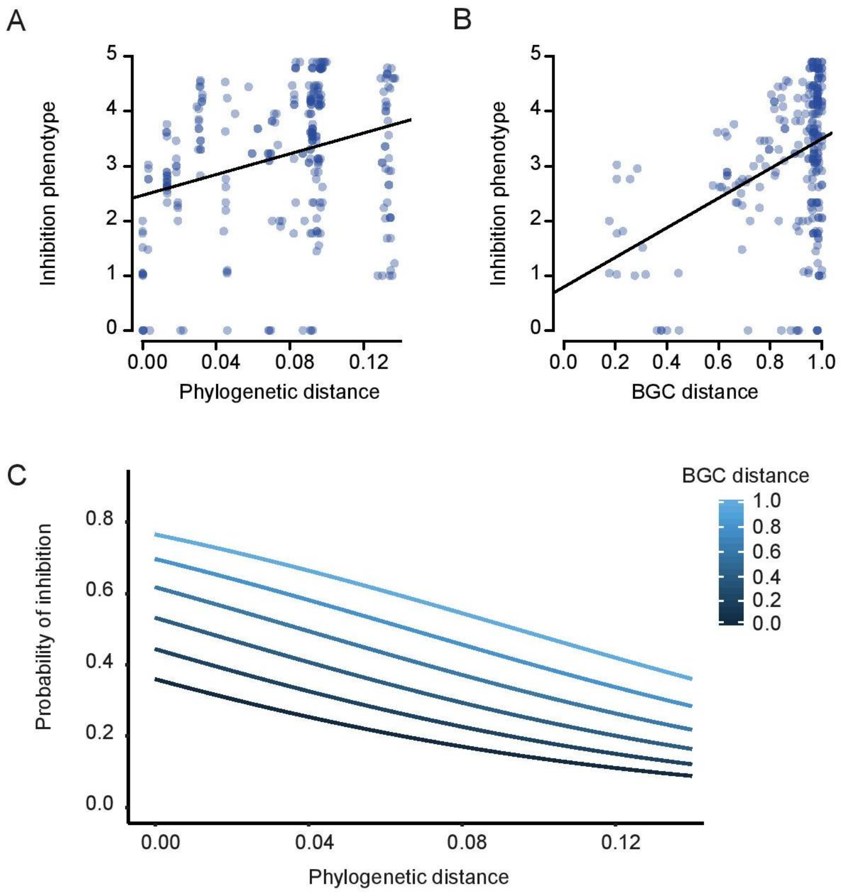


**Fig. S1. Constitutive antagonism under low resource conditions** (A) Correlation between inhibition phenotype dissimilarity and phylogenetic distance (Mantel test, $P < 0.001$, r = 0.30, N = 552) or (B) biosynthetic gene cluster (BGC) distance (Mantel test, $P < 0.001$, r = 0.39, N = 552). (C) Logistic regression between the probability of inhibition and phylogenetic and biosynthetic gene cluster (BGC) distance ($P_{\text{phylogenetic distance}} < 0.001$, $P_{\text{BGC distance}}$ < 0.001, McFadden $R^2 = 0.02$, N = 526).

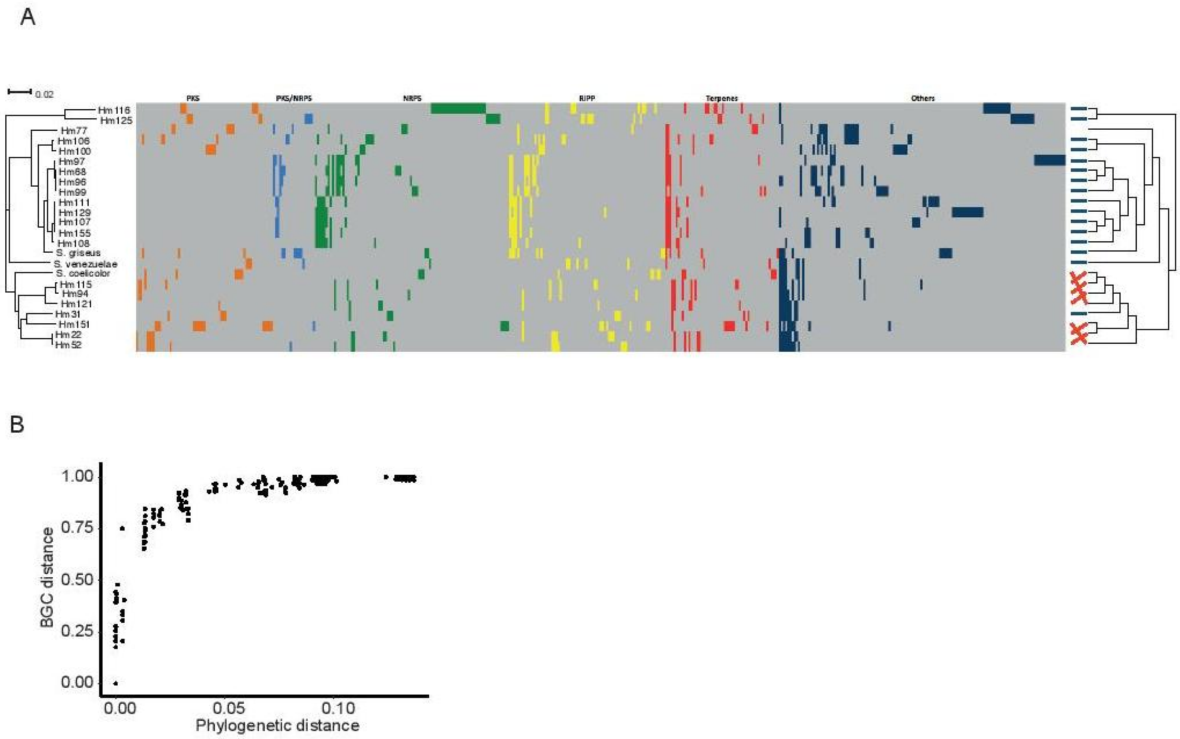


**Fig. S2. Presence/absence of secondary metabolite gene cluster families in all strains.** Biosynthetic gene clusters (BGCs) were identified in all genomes using antiSMASH (22) and BiG-SCAPE (23) was used to cluster the BGCs into BGC families. In the map each column denotes a gene cluster family and each row denotes a strain respective to the phylogeny on the left. The presence of a BGC in the genome belonging to a gene cluster family is indicated by a colour code according to the class: polyketide syntases (PKSs) – orange, polyketide syntases/non-ribosomal peptide synthases hybrids (PKS/NRPS-hybrids) – light blue, non-ribosomal peptide syntases (NRPSs) – dark green, ribosomally synthesized and post-translationally modified peptides (RiPPs) – yellow, terpenes – red, all other identified biosynthetic gene classes – dark blue. The gene cluster families for each class are sorted by abundance from high to low (left to right). Grey indicates the absence of the respective gene cluster family member. The dendrogram on the right (Unweighted Pair Group Method with Arithmetic mean clustering with Jaccard distance) shows the BGC similarity between the strains and is derived from a similarity matrix containing information on the presence/absence of biosynthetic gene cluster families. Blue lines connect strains that are placed similarly in the dendrogram and the phylogeny, while red lines indicate strains that have a different position (B) Correlation between phylogenetic and BGC distance (Mantel test, $P < 0.001$, r = 0.80, N = 552)

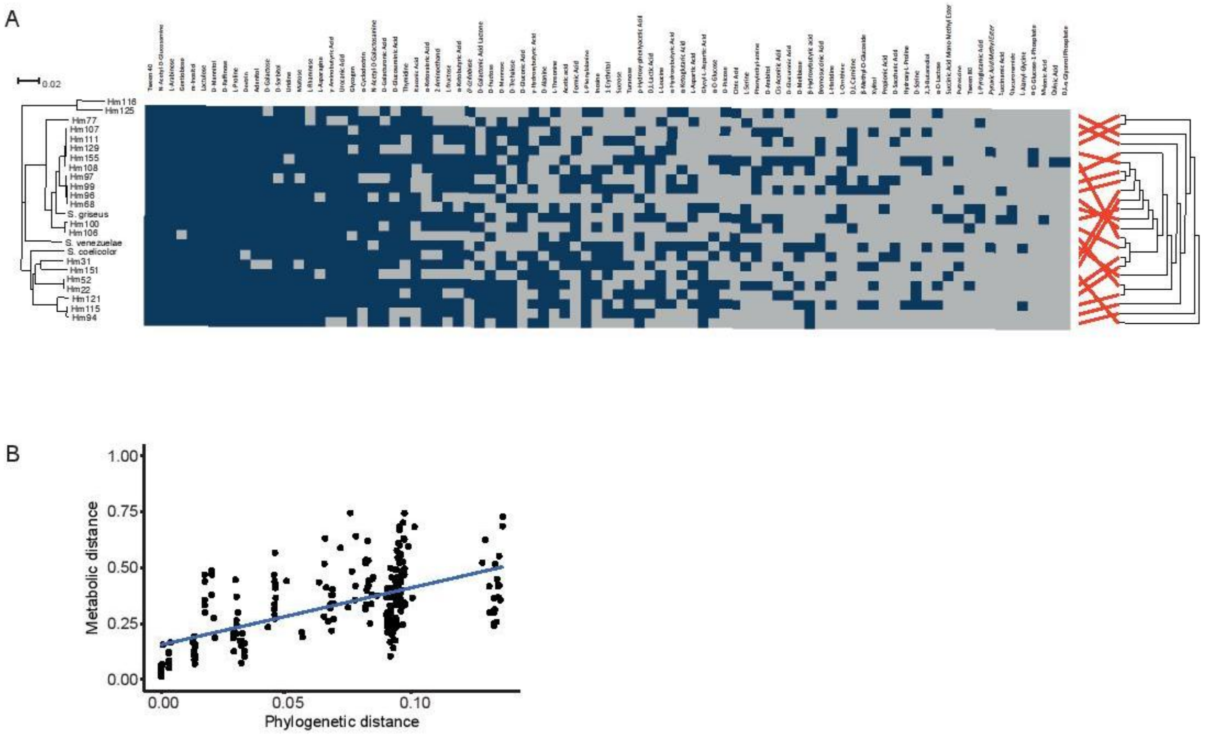


**Fig. S3. Carbon source utilization ability of all strains.** (A) The ability to grow on 95 different carbon sources was assessed using BiOLOG plates. In the map each column denotes a carbon source and each row denotes a strain respective to the phylogeny in Fig. 2A. The ability to grow on a carbon source is indicated in blue, while grey indicates the absence of growth compared to a water control. The dendrogram on the top (UPGMA clustering with Jaccard distance) shows the carbon source utilization similarity between strains and is derived from a similarity matrix containing information on the presence/absence of growth. Strain P (Hm116) showed extremely poor growth on the BiOLOG plates and was excluded from this analysis. (B) Correlation between phylogenetic and metabolic distance (Mantel test, $P < 0.001$, r = 0.60, N = 552).

**Table S1. Strain names corresponding to the identifying letters in the main figures.** All strains designated 'Hm' were isolated from the same soil sample taken from the Himalaya (37). Most of these strains were included in the MBT set described in the same paper and can be found under the corresponding 'MBT' names in other publications.

| **Identifier** | **Strain** |
|---|---|
| A | Hm22 |
| B | Hm31/MBT33 |
| C | Hm52/MBT49 |
| D | Hm68/MBT51 |
| E | Hm77 |
| F | Hm94/MBT53 |
| G | Hm96/MBT54 |
| H | Hm97/MBT55 |
| I | Hm99/MBT56 |
| J | Hm100/MBT57 |
| K | Hm106/MBT58 |
| L | Hm107/MBT59 |
| M | Hm108/MBT60 |
| N | Hm111/MBT61 |
| O | Hm115/MBT62 |
| P | Hm116/MBT63 |
| Q | Hm121/MBT65 |
| R | Hm125/MBT66 |
| S | Hm129/MBT67 |
| T | Hm151/MBT70 |
| U | Hm155/MBT72 |
| V | S. coelicolor A3(2) M145 |
| W | S. venezuelae ATCC 15439 |
| X | S. griseus IFO 13350 |
